# SPRING: a kinetic interface for visualizing high dimensional single-cell expression data

**DOI:** 10.1101/090332

**Authors:** Caleb Weinreb, Samuel Wolock, Allon Klein

## Abstract

**Motivation:** Single-cell gene expression profiling technologies can map the cell states in a tissue or organism. As these technologies become more common, there is a need for computational tools to explore the data they produce. In particular, existing data visualization approaches are imperfect for studying continuous gene expression topologies.

**Results:** Force-directed layouts of k-nearest-neighbor graphs can visualize continuous gene expression topologies in a manner that preserves high-dimensional relationships and allows manually exploration of different stable two-dimensional representations of the same data. We implemented an interactive web-tool to visualize single-cell data using force-directed graph layouts, called SPRING. SPRING reveals more detailed biological relationships than existing approaches when applied to branching gene expression trajectories from hematopoietic progenitor cells. Visualizations from SPRING are also more reproducible than those of stochastic visualization methods such as tSNE, a state-of-the-art tool.

**Availability:** https://kleintools.hms.harvard.edu/tools/spring.html,https://github.com/AllonKleinLab/SPRING/

## 1 Introduction

A recurring challenge in biology is to understand the diversity of molecular states in tissues and organisms. Recent advances in single-cell RNA sequencing (scSeq) (Klein *et al.,* 2015; Trapnell, 2015) have partly addressed this problem, by making it possible to catalogue the expression of every gene in every cell from a given sample with reasonable accuracy. With such data it is now possible to identify the structure of gene expression states, without preconceptions, from population snapshots of single-cells. However, there remains a need for computational tools to explore and visualize the large amount of high-dimensional data generated in scSeq experiments.

When analyzing the distribution of cell states from single-cell gene expression data, two features are of principal interest. First, classifying cells into discrete clusters can give a coarse-grained representation of different phenotypic states (e.g. cell types in a tissue). Second, understanding how cells link together to form continuous topologies can reveal dynamic behaviors during development, differentiation and perturbation response. Currently, the dominant methods for exploring scSeq data are adept at finding clusters in a heterogeneous population, but they tend not to preserve information about how the clusters relate to each other or how they link into continuous topologies in gene expression space. For example, the widely adopted t-distributed Stochastic Neighbor Embedding (tSNE) (Amir *et al.,* 2013; Hinton and Bengio, 2008) is powerful at revealing discrete structure in data, but is not robust with respect to global topology.

To obtain information on continuum processes, graph representations of scSeq data have been used as an alternative to clustering. In a k-nearest neighbor (knn) graph, each cell is a node that extends edges to the k other nodes with most similar gene expression. Several existing approaches for analysis of high-dimensional single-cell data rely internally on a knn graph data structure (Samusik *et al.,* 2016; Xu and Su, 2015; Setty *et al.,* 2016; Bendall et *al.,* 2014; Levine *et al.,* 2015), and one study proposed the use of knn graphs for visualization and data clustering (Islam et *al.,* 2011).

We have found that interactively exploring graph topology, overlaid with gene expression or other annotations, provides a powerful gestalt approach to uncover biological processes emerging from data. Unlike methods that utilize knn graphs for computation, data-enhanced interactive graphs allow intuitive data exploration and can be easily used by biologists without computational training. In addition, interactive graphs can quickly reveal whether observed data structures are robust to graph rearrangements, or whether they reflect an arbitrary distortion of data projected into two dimensions. However, at present there are no publicly available tools for interactive visualization of scSeq data in a graph format.

Here we present a user-friendly web tool called SPRING. To use the tool, users must supply a table of gene expression measurements for single-cells and can optionally upload additional annotations. SPRING builds a knn graph from this data and displays the graph using a force-directed layout algorithm that renders as a real-time simulation in an interactive viewing window.

We include a set of features for open-ended data exploration, including interactive discovery of marker genes; gene expression comparisons between different subpopulations; and selection tools for isolating subpopulations of interest. SPRING is compatible with all major web browsers and does not require technical knowledge to operate.

## 2 Methods

To generate the knn graph, SPRING performs the following transformations to the inputted gene expression matrix. All parameters labeled “X” in this section can be adjusted using an interactive web form.

1. Filter all cells by total read count
2. Normalize so that every cell has the same total reads
3. Filter genes by mean expression and Fano factor
4. Z-score normalize expression values for each gene
5. Perform principal components analysis (PCA), keep the top principal components
6. Compute a distance matrix and output a knn-graph

SPRING is demonstrated in two examples in Figure 1. The underlying data sets are being published in separate research papers, and are available at https://kleintools.hms.harvard.edu/tools/spring.html.

**Figure 1:**
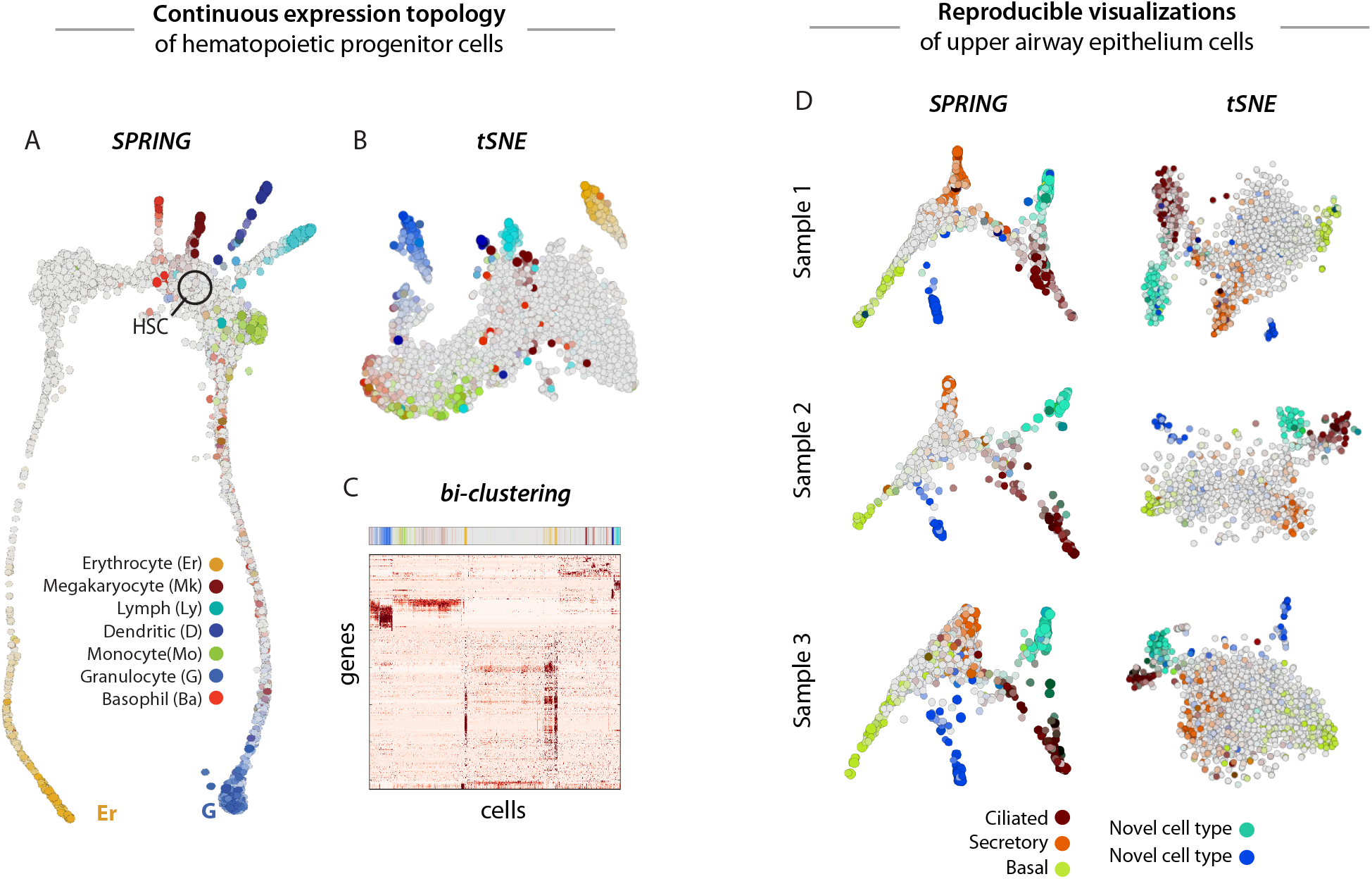
(A) SPRING depicts the dynamic trajectories of hematopoietic progenitor cells as they differentiate from stem cells (HSCs; black circle) into each of seven lineages (colored arms). By contrast, tSNE (B) and bi-clustering (*C*)visualizations of the same data show disconnected clusters of cells, obscuring the underlying continuum process.(*D*)SPRING and tSNE plots of upper airway epithelium cells from three human donors highlight the reproducibility of SPING visualizations. Cells in (A-D) are colored by sum-Z-score of marker genes for each respective cell type. Cell type markers for figs 1A-C are provided as a supplementary file in SPRING format.

## 3 Advantages over existing methods

### 3.1 Continuous expression topologies

In contrast to methods such as tSNE and bi-clustering, SPRING captures the long-distance relationships between cells and can therefore visualize continuous expression topologies. For example, SPRING accurately maps the branching topology of hematopoietic progenitor cells as they first choose between and then differentiate along seven lineages (Figure 1A). In contrast, tSNE (Figure 1B) and bi-clustering (Figure 1C) do not capture the correct expression topology for these cells.

### 3.2 Graph invariance

One drawback of tSNE - the state-of-the-art algorithm for visualizing high dimensional scSeq data - is that it is stochastic and therefore not perfectly reproducible. We have observed highly variable results when running tSNE on different biological replicates of the same tissue or even multiple runs on the same dataset. By contrast, graph construction in SPRING is non-stochastic and therefore yields consistent topologies between runs and replicates. In addition, manual interaction with the kinetic SPRING interface allows users to bring plots from separate replicates into register with one other (Figure 1D).

## 4 Conclusion

Single-cell gene expression profiling is becoming a common tool to dissect cellular heterogeneity and characterize dynamic processes such as differentiation. Interactive visualization tools can help researchers exploit this data more fully. Our easy-to-use web tool, SPRING, provides a simple interface for open-ended investigation of gene expression topology. In contrast to existing methods such as tSNE, SPRING accurately portrays continuum processes and generates visualizations that are highly reproducible.

## Acknowledgements

We thank James Briggs and Virginia Savova for helpful comments and suggestions.

